# Reward modulates behaviour and neural responses in motor cortex during action observation

**DOI:** 10.1101/868083

**Authors:** Andreea Loredana Cretu, Rea Lehner, Rafael Polania, Nicole Wenderoth

## Abstract

Transcranial magnetic stimulation (TMS) studies demonstrated that observing the actions of other individuals leads to action-specific facilitation of primary motor cortex (M1) (i.e., “motor resonance”). Motor resonance is modulated by contextual information accompanying others’ actions, however, it is currently unknown whether action value influences behavioural and physiological outcomes during action observation in humans. Here we tested whether response times (RT) and muscle-specific changes of M1 excitability are modulated by the value an observer assigns to the action executed by another agent and whether this effect can be distinguished from attentional engagement. We show that observing highly-valued actions leads to a significant decrease in RT variability and a significant strengthening of action-specific neural representations in M1. This “sharpening” of behavioural and neural responses was observed over and beyond a control task requiring similar attentional engagement but did not include any rewards. Our finding that reward influences action specific representations in human M1 even if no motor response is required is new, suggesting that reward influences the transformation of action stimuli from the perceptual to the motor domain. We suggest that premotor areas are important for mediating the observed effect, most likely by optimizing grasp-specific PMv-M1 interactions which cause muscular facilitation patterns in M1 to be more distinct for rewarded actions.

## 1. Introduction

Observing the movements performed by another individual automatically triggers changes in corticospinal excitability measured in the primary motor cortex (M1), a phenomenon called “motor resonance”. This modulation of M1 activity is thought to reflect the influence of upstream parietal and premotor areas which form the mirror neuron system (MNS) (for a review, see Naish et al., 2014). It has been shown that M1 activity during action observation depends not only on the observed kinematics but is additionally modulated by higher-order information, such as the context in which the action is taking place (Amoruso & Urgesi, 2016), characteristics of the agent performing the action (Bunday, Lemon, Kilner, Davare, & Orban, 2016) or moral connotations of observed actions (Craighero & Mele, 2018). Yet, the contribution of action value (i.e., reward associated with observed actions) to M1 motor resonance in humans has remained unclear.

The only direct evidence for the influence of reward on the MNS activity comes from a recent study in primates (Caggiano et al., 2012). Caggiano and colleagues identified mirror neurons in area F5 of the monkey premotor cortex that modulated their firing rates depending on the subjective value associated with the observed actions. In particular, they showed that neural firing of grasp-specific mirror neurons was modulated by whether the grasped object was rewarding or non-rewarding. While most neurons fired when the subjective reward value of the object grasped by another agent was high, some had a preference for less-relished or no rewards. Thus, the observer’s subjective value associated with the to-be-grasped object modulated how the observed action was represented by mirror neuron activity, an effect that was observed over and above potential attention-related responses.

In humans, the existence of a similar interaction between subjective value representations and motor resonance (i.e. brain activity reflecting observed motor actions), can only be drawn on the basis of indirect evidence. Previous studies proposed that electrophysiological markers of motor resonance (i.e., decreased power in mu band measured with electroencephalography (EEG)) are modulated by the observation of rewarding stimuli (Brown, Gonzalez-Liencres, Tas, & Brüne, 2016; Brown, Wiersema, Pourtois, & Brüne, 2013; Trilla Gros, Panasiti, & Chakrabarti, 2015). In particular, they found more pronounced mu suppression (interpreted as stronger motor resonance) when observing rewarding actions which suggests that the higher subjective value of an observed action, the stronger is the response of the observers’ sensorimotor system. However, it has been argued that mu suppression is only a reliable indicator of MNS activity when attentional engagement is tightly controlled (Hobson, Bishop, & Hobson, 2016) such that the hypothesis of whether the subjective value of an observed action modulates M1 activity over and beyond attentional effects remains untested in humans.

Here we addressed this question via a behavioural and a TMS experiment using a well-established action observation paradigm (Cretu, Ruddy, Germann, & Wenderoth, 2019; de Beukelaar, Alaerts, Swinnen, & Wenderoth, 2016) where participants observed either a precision grip or a whole hand grip. We contrasted different experimental conditions where (i) the different grip types were associated with high or low reward (reward condition); or (ii) had to be counted or ignored (attentional control).

## Materials and Methods

In two separate experiments, we examined whether response times (experiment 1) and muscle-specific motor resonance (experiment 2) are modulated by the value associated with an observed action. Both involved participants viewing grasping movements presented on a computer screen.

### 1.1. Movement stimuli

In both experiments, participants observed either a precision-grip (PG) or a whole-hand grip (WHG). These actions were presented in an implied motion paradigm (still shots of two different grasping movements; **Figure 1**). We chose to show static images presented in succession to give the impression of movement rather than complete action videos in order to have more control over the timing of the sequence of events. Stimuli were created by showing a series of three still shots from videos of a hand reaching, grasping and lifting a round jar (9 cm diameter, 14.5 cm height, 1.5 kg weight). These still shots were chosen from videos that were used previously (Cretu et al., 2019; de Beukelaar et al., 2016). In WHG all fingers were used to grasp the lid of the jar, whereas for the PG, only the thumb and index finger grasped a small knob mounted on the top of the jar. Each trial started with a fixation cross which stayed on the screen for a random duration between 3.5 and 4.5 seconds followed by the reach, grasp and lift pictures – each remaining visible for 0.6 seconds. In the reach picture, the jar was visible in the middle of the screen together with the right hand of an actor positioned in the upper right corner as a clenched fist. Importantly, both WHG and PG condition were identical at this phase. The grip type only became apparent in the second and third pictures which showed the hand grasping the jar and lifting it approximately 15 cm above the surface. The occurrence of WHG and PG trials was randomized to avoid anticipation on the type of grasp.

**Figure 1.**
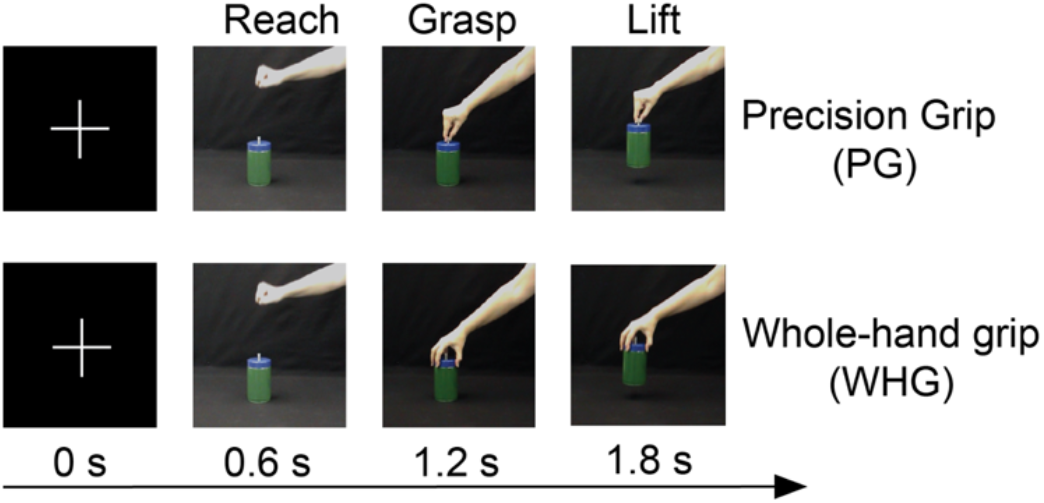
Movement stimuli. Schematic representation of a precision grip (PG) and a whole-hand grip (WHG) trial structure. A fixation cross was presented for a random duration (3.5 – 4.5 seconds) followed by the reach, grasp and lift still shots (0.6 seconds each). During the experiment, PG and WHG trials were randomly intermixed.

Depending on the experiment, we either recorded response-times (RTs; experiment 1) or motor-evoked potentials (MEPs; experiment 2) during the grasping snapshot.

#### Reward condition and counting control

Both experiments employed two main conditions: a **reward** condition and a **counting** control condition. Additionally, in each block, one of the two grasp types was task-relevant (i.e., the **target**) while the other was task-irrelevant (i.e., the **non-target**). In the reward condition, subjects had to count the accumulated monetary reward directly associated with the target, while in the counting condition, they had to count how many times they saw the target.

### 1.2. Experiment 1: behavioral effects of reward during action observation

#### 1.2.1. Participants

49 volunteers were recruited for this experiment. Of these, two were excluded from the final analyses because of their high rate of wrong button presses (i.e., button presses in more than 10% non-target trials). Thus, the final sample for this study included 47 participants (25 women, mean age ± sd 27.25 ± 6.73 years old, right-handed). All experimental protocols were approved by the research ethics committee of the canton of Zurich (no. 2017−01001). Participants gave informed consent to the study, were paid for participation and received a random cash bonus which depended on their performance during the experiment (max. 20 Swiss francs; for more information see the *Behavioral paradigm* section).

#### 1.2.2. Behavioral paradigm

In this experiment, we assessed the behavioral effects of reward during action observation by analyzing response times. Each trial always started with a fixation cross which stayed on the screen for a random duration between 3.5 and 4.5 seconds followed by the reach, grasp and lift pictures – each remaining visible for 0.6 seconds. The occurrence of WHG and PG trials was randomized to avoid anticipation on the type of grasp.

Participants had to perform 18 blocks each consisting of 16 trials (on average: 50% PG and 50% WHG). Prior to each block they were instructed to attend to one of the grasp types (i.e., the target) while the other was task-irrelevant (i.e., the non-target). Importantly, one of three different values could be assigned to the target action: 2 Swiss francs (i.e., high reward blocks, n=6), 0.02 Swiss francs (i.e., low reward blocks, n=6) or 0 Swiss francs (i.e., counting control blocks, n=6), as it can be seen in **Figure 2** below. Participants pressed a button whenever they saw the target action (PG or WHG, depending on the block). In reward blocks, subjects had to additionally count the accumulated reward associated with the target grasp. After the button press was recorded, the respective monetary reward was shown as a coin appearing during the lifting snapshot to remind participants that this is the task-relevant target grasp and whether they could earn a low or high reward. After each block, the accumulated sum of reward was written down and the participants were informed that only the *correct* sum is considered for renumeration (see below). The counting control blocks followed a similar structure. The main difference was that participants had to simply count how many times they saw the target grasp. After each block, the count of the target action was written down and the participants were informed that only the *correct* count is considered for renumeration (see below). All tasks required the same simple button press (i.e., spacebar) to ensure that any resulting differences in response time across these blocks were dependent only on the reward expectancy (i.e., high vs. low reward vs. counting), and not driven by differences in task difficulty. Thus, depending on the block instructions, we either investigated reward effects (target action is rewarded) or attentional effects (target action is counted).

**Figure 2.**
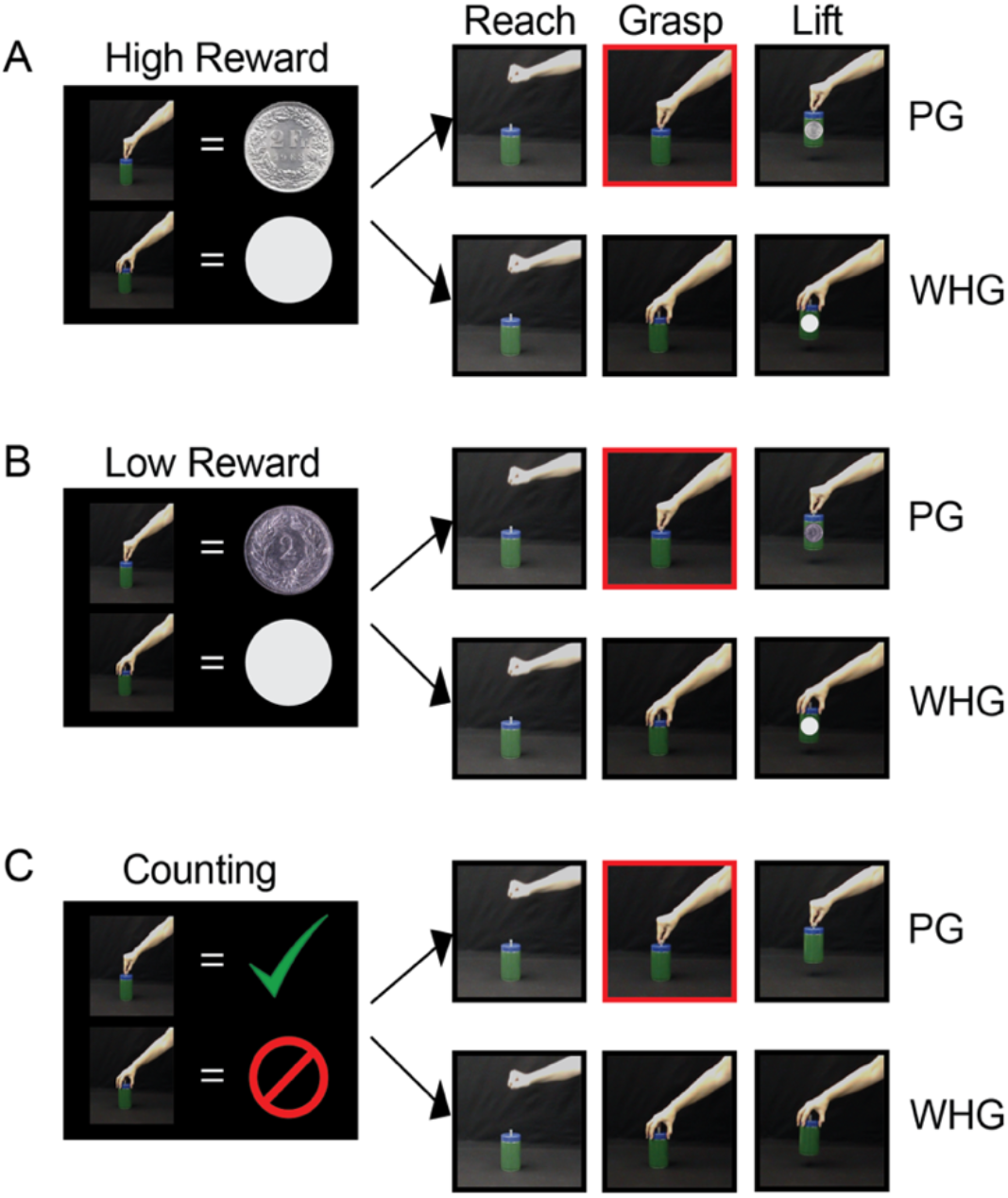
Illustration of behavioral task instructions (left part of each panel) and trial structure (rigt part of each panel) All examples depict blocks in which precision grip (PG) is the target and whole-hand grip (WHG) is the non-target action. The red frame indicates the snapshot during which the response times were recorded. (A) High reward block: 2 Swiss francs associated with the target. (B) Low reward block: 0.02 Swiss francs associated with the target. (C) Counting attentional block: no monetary reward associated with the target.

Prior to the experiment, participants were informed about our renumeration scheme which incentivised them to have a good performance throughout the whole duration of the experiment. In particular, participants were instructed that only one block will be picked randomly and remunerated at the end of the experiment. If a reward block was chosen, the amount of money depended on the total sum accumulated in the block but only when this sum was reported correctly. If a counting block was chosen, participants were informed that although no value was directly associated with the target grasp, a fixed reward was offered if the reported count was correct. This payoff structure ensured that subjects were motivated to attend to the target grasp independent of the block type.

Feedback was never given on a trial-by-trial basis since this might lead to reduced task-performance (Kluger & Denisi, 1996) and unwanted trial-by-trial dynamics. However, feedback was given at the end of each block. Subjects only had to press a button for the target grasps. In total, 24 RTs were recorded for each of six experimental conditions, i.e. 2 grip types (WHG, PG) x 3 tasks (low reward, high reward counting), resulting in a total of 144 RTs for each subject.

#### 1.2.3. Behavioral analysis

Response time (RT) was the primary outcome variable of this experiment and was defined as the time between grasp snapshot onset and the button press. Occurrences of unusually quick responses (<200 ms which is faster than possible to recognize the target grasp and perform a motor response, as suggested by Naish et al., 2014) were removed from the analyses (0.76 % of all trials). Subjects only had a short time window of 600ms to respond to the target grasp and if they were too slow to respond in that time window, it counted as a miss (0.2 % of all trials).

An Ex-Gaussian model was fitted to the remaining RT data for each condition and subject and three model parameters (mu, sigma, and tau) were estimated using the Matlab toolbox exgauss (Zandbelt, 2014). The advantage of ex-gaussian fitting is that the positive skewness of response time distributions is considered by combining a normal Gaussian distribution with an exponential component. The mu and sigma parameters reflect the mean and standard deviation (sd) of the gaussian component of the RT distribution while tau describes the exponential distribution (Lacouture, 2008). Chi-square tests confirmed that the ex-Gaussian model was a good fit of the individual data of each condition for 46 out of the 47 participants. Next, ex-Gaussian parameters of all participants were subjected to a boxplot method to identify and remove extreme data points (n=5) that exceeded Q1-1.5*IQR or Q3+1.5*IQR respectively with Q1 being the first quartile, Q3 the third quartile and IQR the interquartile range computed over the whole data set (Tukey, 1977).

Counting errors (i.e., wrong calculation of the accumulated reward or wrong counting) as well as incorrect, early and late button presses (i.e., button presses during non-target grasps, during reach or lift respectively) were inspected to ensure that cognitive demand was comparable across the three experimental tasks (i.e., low, high reward or counting).

#### 1.2.4. Statistical analysis

To investigate the effects of observed action value (low, high reward, and count) on behavioral performance, ex-Gaussian parameters (mu, sigma, tau) were analyzed using repeated-measures analysis of variance (RM-ANOVA) in JASP (Version 0.8.5.1, https://jasp-stats.org/). Error rates were analysed in the same manner.

Estimates of the effect sizes are reported in the tables using partial eta squared (small 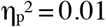, medium 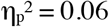, large 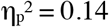; Lakens & Bakker, 2013). The significance threshold alpha=.05 was chosen for all statistical tests and FDR corrections for multiple comparisons were applied when necessary. Results are reported in the figures as mean ± standard error of the mean (SEM). Additionally, we added estimation plots to depict Cohen’s d effect sizes (small d = 0.20-0.49, medium d = 0.50–0.80, large d > 0.80; Cohen, 1988) and their confidence intervals in form of bootstrap sampling distributions using open-source code (*http://www.estimationstats.com;* Ho, Tumkaya, Aryal, Choi, & Claridge-Chang, 2019).

### 1.3. Experiment 2: Transcranial Magnetic Stimulation Experiment

#### 1.3.1. Participants

34 volunteers were recruited for a transcranial magnetic stimulation (TMS) experiment (19 women, mean age ± sd 25.8 ± 4.8 years old). All subjects were right-handed, as assessed with the Edinburgh Handedness Questionnaire (Oldfield, 1971), naive about the purpose of the experiment and had normal or corrected-to-normal vision. All participants were screened for potential contraindications to brain stimulation (Rossi et al., 2009) and written informed consent was obtained before starting the task. Subjects were paid for participation but additionally received a random cash bonus which depended on their performance during the experiment (max. 20 Swiss francs; more information in the *Behavioral paradigm* section). All experiments were approved by the research ethics committee of the canton of Zurich (no. 2017-01001).

#### 1.3.2. Behavioral paradigm

The goal of this experiment was to test the effects of reward on action-specific responses elicited during action observation. The task design was the same as in Experiment 1, except for the following two changes. First, participants were instructed to be as relaxed as possible during the task and to silently count either the accumulated reward or the number of target grasps (i.e., no button press required during the grasp presentation). Second, in the reward blocks, we showed either 1 or 2 Swiss francs coins during the lifting phase randomly interleaved within a block which they had to sum up and report at the end of each block (**Figure 3**). We chose two different reward levels to ensure that subjects focused on the value of the action and did not simply count the target grasps. For non-target trials, a coin-like circle appeared in the same location and at the same time point.

**Figure 3.**
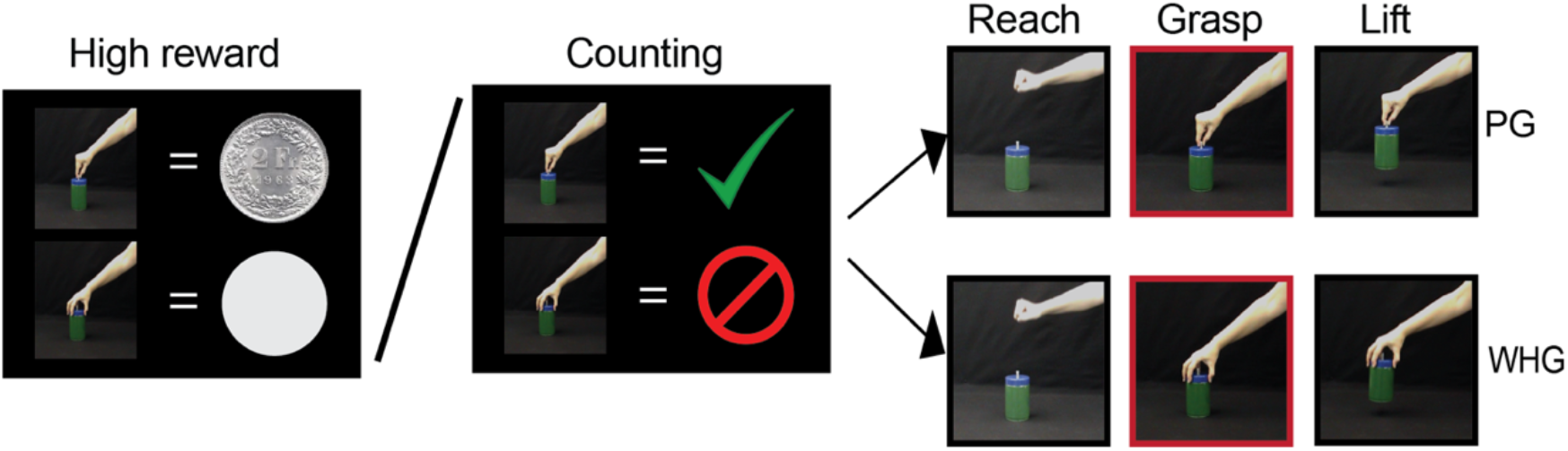
Illustration of TMS task instructions and trial structure. Schematic representation of the TMS task instructions and examples of precision grip (PG) and whole-hand grip (WHG) trials. Instructions depict a high reward and a counting block in which PG is the target and WHG is the non-target action. The red frame indicates the snapshot in which the TMS pulses were sent and motor - evoked potentials were recorded.

The payoff structure was similar to the one presented in Experiment 1. Namely, one random block was picked and remunerated at the end of the experiment, if subjects performed the task correctly. The amount of money corresponded to the sum accumulated in the respective block (reward condition) or was a fixed sum which was given for good performance (i.e., correct counting; count condition). TMS single-pulses were sent after 250 ms from the presentation of the grasp snapshot because previous electrophysiological and TMS research showed that motor resonance can be reliably observed around 250 ms after action onset (Barchiesi & Cattaneo, 2013; Cavallo, Heyes, Becchio, Bird, & Catmur, 2014; Ubaldi, Barchiesi, & Cattaneo, 2015; C. Urgesi, Moro, Candidi, & Aglioti, 2006; Cosimo Urgesi et al., 2010).

Here, for each of the eight conditions [2 grip types (WHG, PG) x 2 target types (target, non-target) x 2 tasks (reward, counting)], 24 MEPs were recorded, resulting in a total of 192 MEPs for each subject and each muscle (FDI, ADM).

#### 1.3.3. Electromyographic recordings and TMS procedure

Corticomotor excitability was measured using TMS and motor-evoked potentials (MEPs) were simultaneously recorded from the First Dorsal Interosseous (FDI) and Abductor Digiti Minimi (ADM) of the right-hand using surface electromyography (EMG; Delsys Bagnoli DE-2.1). EMG data were recorded using Signal Software (Version 5.07, Cambridge Electronic Design, UK), sampled at 5000 Hz (CED Power 1401; Cambridge Electronic Design, UK), amplified, filtered with a band-pass (30–1000 Hz) and a notch (50Hz) filter, and stored on a PC for off-line analysis. TMS was performed by means of a 70 mm figure of eight coil connected to a Magstim200 stimulator (Magstim, Whitland, UK). Since we wanted to make sure that MEPs are consistently elicited in both muscles, we defined the optimal scalp position (hotspot) as the position from which stable responses were simultaneously evoked in both the right FDI and ADM muscles. The resting motor threshold (rMT) was defined as the lowest stimulus intensity evoking MEPs in the right FDI and ADM with an amplitude of at least 50 μV in 5 out of 10 consecutive stimuli (Rossini et al., 2015). Subjects’ rMT, expressed as a percentage of the maximum stimulator output, was on average 45 % (range: 34% - 57%). TMS triggering was controlled using Matlab 2013b (The MathWorks, Inc., Natick, USA) in combination with a Teensy 3.1 microcontroller. To reliably record MEPs in both muscles, the stimulation intensity used was 130%rMT as described previously (Alaerts, de Beukelaar, Swinnen, & Wenderoth, 2012; Alaerts, Heremans, Swinnen, & Wenderoth, 2009).

From the EMG data, peak-to-peak amplitudes of the MEPs were calculated using custom-made Matlab scripts (Matlab 2015, Mathworks, Natick, MA, USA). Data preprocessing was done using the same criteria presented previously (Cretu et al., 2019). Shortly, the root mean square (rms) of background EMG in the period 110 to 10ms before the TMS pulses was calculated and MEPs with background rms higher than 10 μV were excluded. Additionally, the mean and standard deviation of the background EMG was calculated for each subject and over all trials and MEPs with background EMG exceeding 2.5 standard deviations above the mean were removed. Finally, MEPs with a peak-to-peak amplitude smaller than 50 μV were also eliminated from further analyses. Following these criteria, a total of 3.88% of all trials were removed from the data. For the remaining MEPs, the mean peak–to-peak amplitude was calculated separately for each muscle, grasp type, target and task condition.

#### 1.3.4. TMS analysis

Both action execution and action observation studies found that the FDI versus ADM muscle are preferentially activated in PG and WHG, respectively (e.g.: Cavallo, Bucchioni, Castiello, & Becchio, 2013; Lemon, Johansson, & Westling, 1995; Sartori, Bucchioni, & Castiello, 2012). In order to quantify such grasp specific changes in M1 we determined an index of MEP modulation. For each muscle (i.e., FDI and ADM) we calculated whether observing a whole-hand grip versus a precision-grip modulated the MEP amplitude:

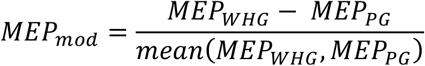

Action-specific motor resonance is signalled by positive MEP_mod_ values for the ADM, indicating a stronger facilitation during WHG than PG observation, and negative MEP_mod_ values for the FDI muscle, indicating a stronger facilitation during PG than WHG observation. The average MEP amplitudes for each condition and muscle are presented in supplementary table 1.

#### 1.3.5. Statistical analysis

*MEP_mod_* was calculated for each subject, muscle and condition and subjected to RM-ANOVA as dependent variable. We employed muscle (FDI, ADM), target (target, non-target) and task (reward, count) as within-subject factors. As a follow-up analysis, we performed one sample t-tests to determine whether MEP_mod_ is larger than 0 for ADM and smaller than 0 for FDI as predicted for an action-specific modulation. Since we had a strong a-priori hypothesis, these tests were one-sided.

## 2. Results

### 2.1. Experiment 1: High rewards lead to decreased response time variability

When comparing the effects of high reward, low reward and counting on the ex-Gaussian model parameters, we found a significant difference in sigma (p = .018; η^2^ = .094, Table 1 and **Figure 4**), representing the standard deviation of the Gaussian component which is related to intra-subject RT variability. Post-hoc analyses revealed that the average sigma in the high reward condition was significantly smaller than in the counting (p_FDR_ = .011, Cohen’s d = .576) and the low reward condition (p_FDR_ = .013, Cohen’s d = .542). No difference was found between sigma in the low reward and counting condition (p_FDR_ = .992). This indicates that actions associated with a high reward were more precise (i.e., less variable) compared to low or no monetary reward.

**Table 1.**
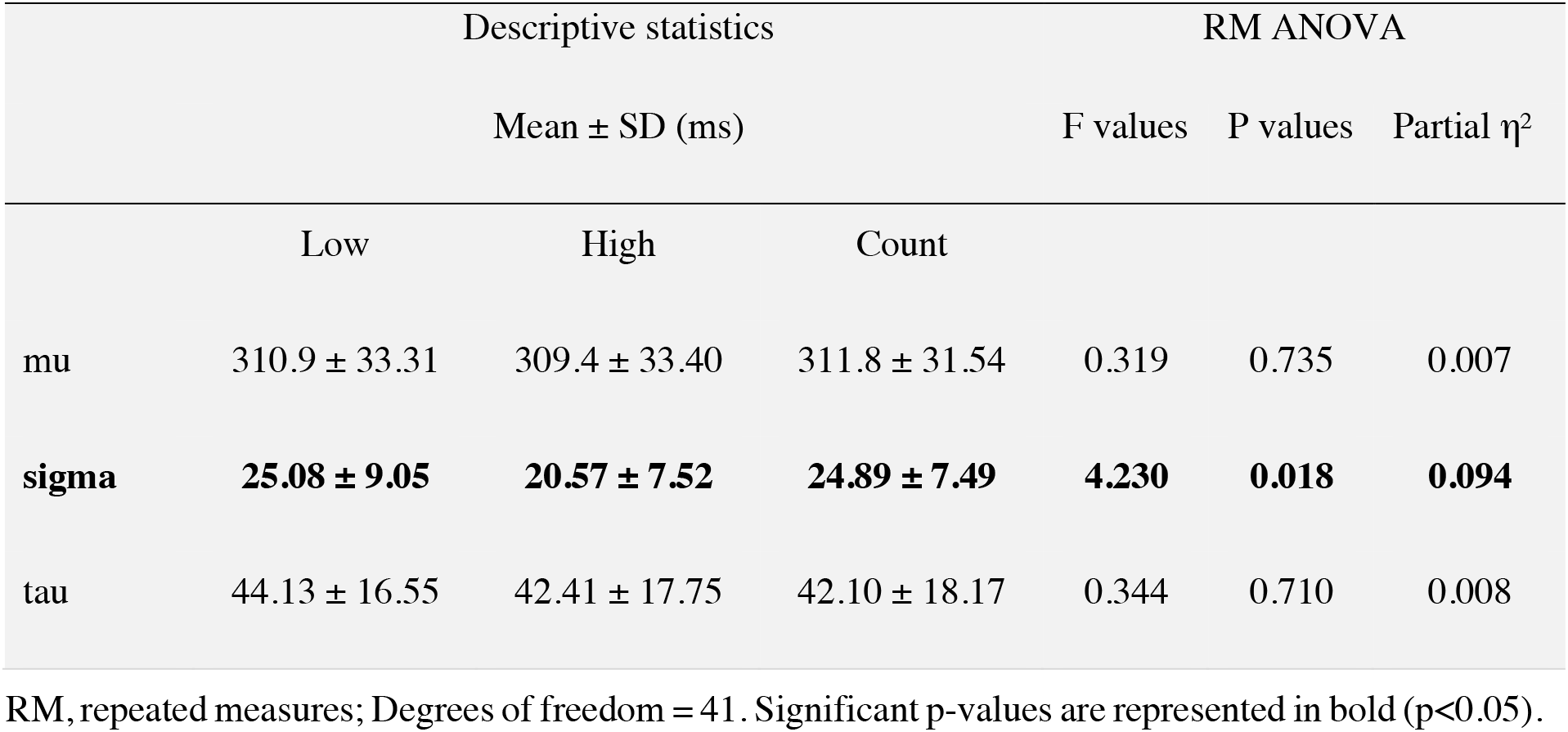
Descriptive and RM ANOVA results for the RT distribution parameters.

**Figure 4.**
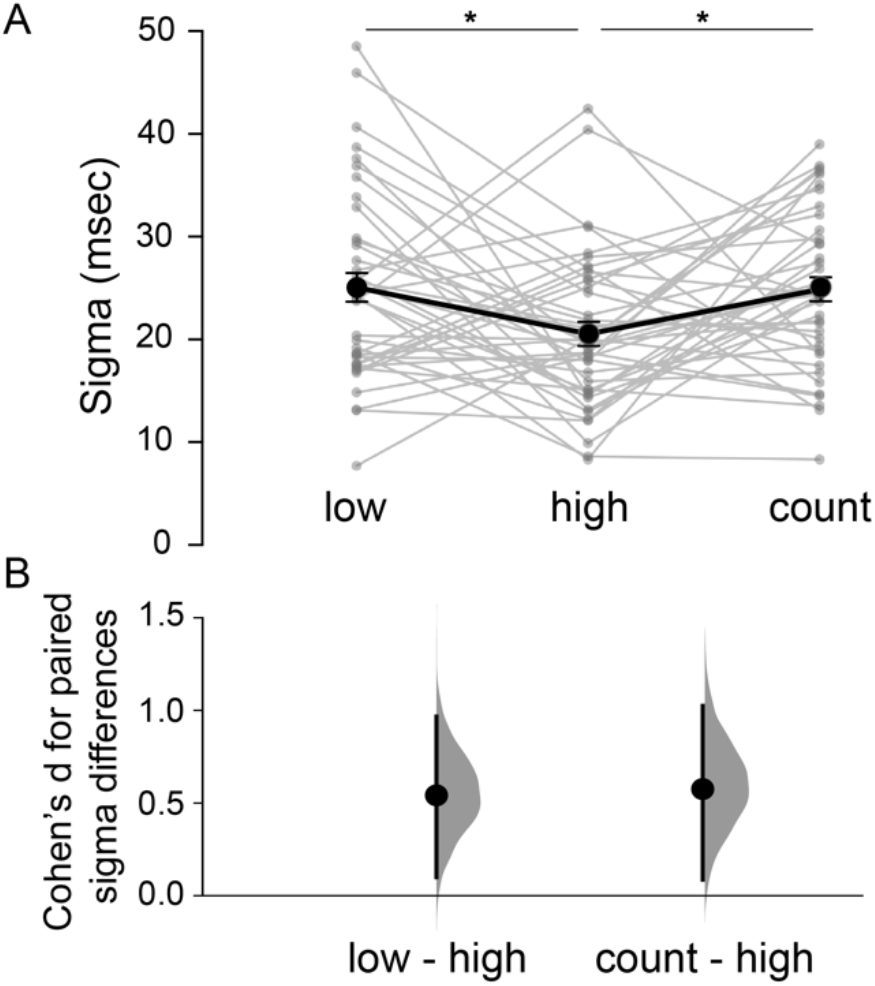
Behavioral results (N = 42) A) Raw sigma values in function of task instructions (i.e., low reward, high reward, counting; n = 42). B) Effect sizes (Cohen’s d) for significant paired sigma differences are depicted as dots. Filled curves depict the resampled distribution of the sigma differences, given the observed data, and error bars depict 95% confidence intervals. *P < 0.05 [false discovery rate (FDR)-corrected t-tests].

No reward versus counting differences were found for the gaussian mean mu (p = .735) or the exponential distribution tau (p =.710). Participants needed approximately 310 ms (Table 1; mu results) to respond to target actions, independent of task instructions. These results suggest that task instructions affect the variability of the responses (i.e., sigma) but not the motor or decisional speed (represented by mu and tau) of the participants.

Furthermore, the number of errors executed within low, high or count blocks were not significantly different (all p >= .37) showing that behavioral performance was unaffected by the task instructions. This suggests that the load on working memory resources was similar across the different conditions.

Results of the descriptive statistics, as well as the repeated measures ANOVA performed on the error rates are presented in Table 2.

**Table 2.** Results of descriptive and RM ANOVA analysis performed with error rates.

|  | Descriptive statistics |  |  | RM ANOVA |  |  |
| --- | --- | --- | --- | --- | --- | --- |
| | Mean ± SD (ms) | | | F values | P values | Partial $\eta^2$ |
|  | Low | High | Count |  |  |  |
| errors counting blocks | 0.40 ± 0.79 | 0.29 ± 0.65 | 0.36 ± 0.48 | 0.482 | 0.592 | 0.010 |
| early presses | 0 | 0.021 ± 0.14 | 0 | 1 | 0.372 | 0.021 |
| late presses | 0.62 ± 1.15 | 0.59 ± 1.13 | 0.78 ± 1.14 | 0.481 | 0.620 | 0.010 |
| wrong presses | 0.62 ± 0.87 | 0.51 ± 1.08 | 0.47 ± 1.02 | 0.310 | 0.734 | 0.007 |

RM, repeated measures; Degrees of freedom = 46. Significant p-values are represented in bold ( $p < 0.05$ ).

### 2.2. Experiment 2: Reward driven sharpening of muscle-specific M1 excitability

We probed the action-specific changes in M1 corticomotor excitability in FDI and ADM muscles during the observation of target (i.e., task-relevant) or non-target (i.e.., task irrelevant) grasping actions embedded in a rewarding (i.e., reward task) or an attentional context (i.e., counting task).

Our results indicate that observing a whole-hand grip versus a precision-grip lead to differential modulation of MEP amplitude in the two muscles (i.e., on average we found MEP_mod_ < 0 for FDI and MEP_mod_ > 0 for ADM; main effect of muscle p<.001, η^2^ = 0.423). Furthermore, this action-specific corticomotor excitability was stronger when observed grasping actions were rewarded versus counted (muscle x task p = .047, η2 = 0.114), as well as when observed grasps was the target action (muscle x target p = .014, η^2^ = 0.17), as can be seen in **Figure 5** and Table 3 (also see Supplementary Figure 2).

**Figure 5.**
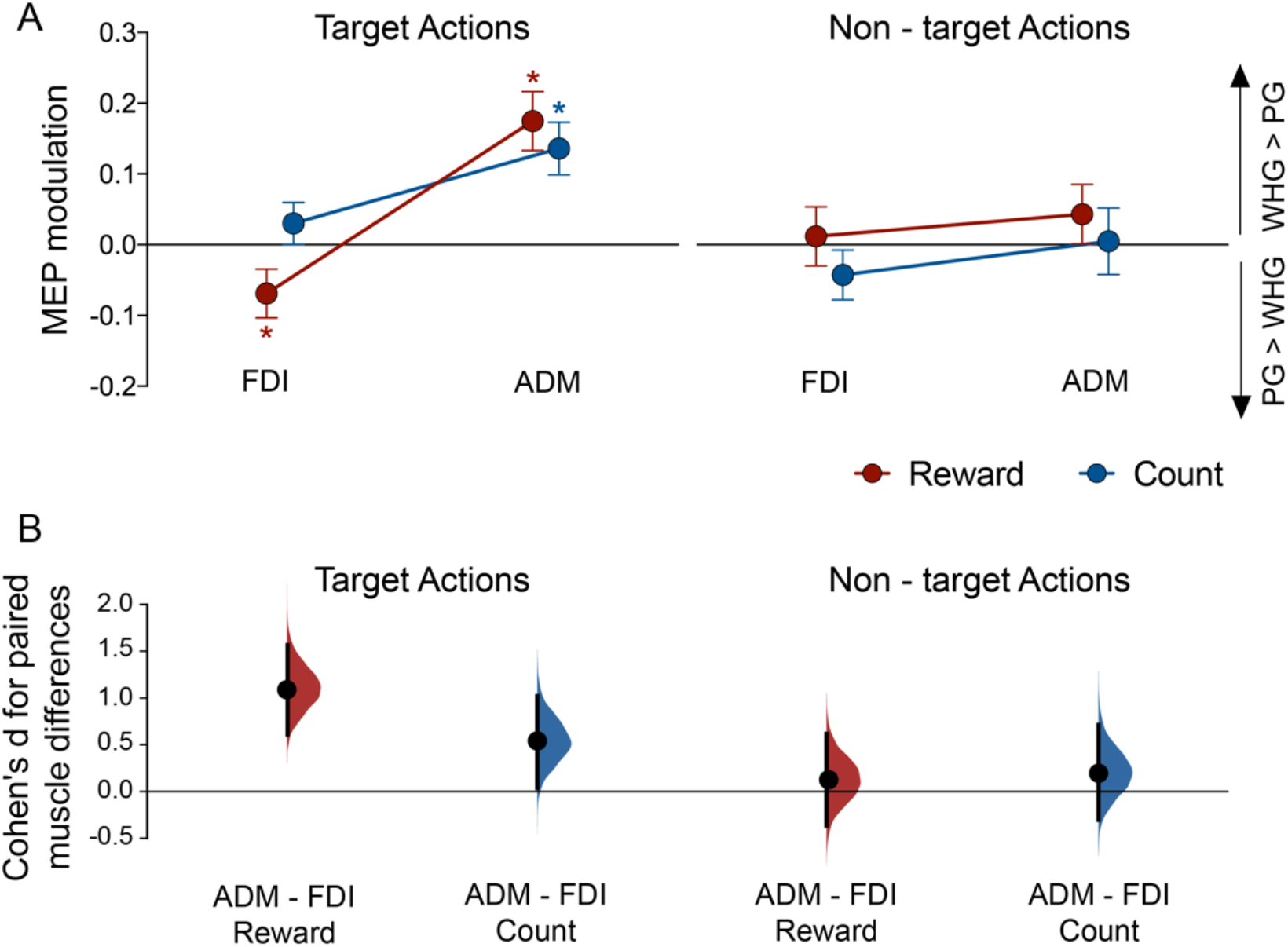
Results of TMS experiment (n = 34). A) Motor-evoked potential (MEP) modulation during the observation of videos depicting a whole hand grip (WHG) or a precision grip (PG) either in a reward (red lines) or counting (blue lines) task. Results are presented separately for each target type and muscle [first dorsal interosseous (FDI) and abductor digiti minimi (ADM)]. Values >0 indicate higher facilitation during WHG vs. PG observation, whereas values <0 indicate increased facilitation during PG vs. WHG observation. B) The action-specificity effect (i.e., ADM – FDI) in each condition is plotted as a bootstrap sampling distribution with Cohen’s d effect sizes depicted by the black dots and the 95% confidence intervals indicated by the vertical error bars. *P < 0.05 [false discovery rate (FDR)-corrected 1-sided t-tests].

**Table 3.** RM ANOVA results performed with the MEP_mod_ data (N = 34)

| | F values | P values | Partial $\eta^2$ |
| --- | --- | --- | --- |
| *Main Effects* |  |  |  |
| **Muscle (FDI, ADM)** | **24.158** | **<0.001** | **0.423** |
| Task (reward, counting) | 0.165 | 0.688 | 0.005 |
| Target (target, non-target) | 3.59 | 0.067 | 0.098 |
| *Interactions* |  |  |  |
| **Muscle x Task** | **4.244** | **0.047** | **0.114** |
| **Muscle x Target** | **6.753** | **0.014** | **0.17** |
| Task x Target | 0.838 | 0.367 | 0.025 |
| Muscle x Task x Target | 1.722 | 0.199 | 0.05 |
| *Target action* |  |  |  |
| **Muscle (FDI, ADM)** | **27.27** | **<0.001** | **0.452** |
| Task (reward, counting) | 0.539 | 0.468 | 0.016 |
| **Muscle x Task** | **5.55** | **0.025** | **0.144** |
| *Non-target action* |  |  |  |
| Muscle (FDI, ADM) | 1.293 | 0.264 | 0.038 |
| Task (reward, counting) | 0.823 | 0.371 | 0.024 |
| Muscle x Task | 0.053 | 0.820 | 0.002 |

ADM, abductor digiti minimi; FDI, first dorsal interosseous; RM, repeated measures; Degrees of freedom = 33. Significant p-values are represented in bold ( $p < 0.05$ ).

To better understand the effects of action-value on motor resonance, we calculated a reduced statistical model, separately for the target and non-target actions. Results showed that during the observation of target grasps, action-specific modulation of MEPs is present and influenced by the reward versus counting instructions (muscle p < 0.001, η^2^ = 0.452; muscle x task p = .025, η^2^ = 0.144). Monetary rewards caused a significantly stronger change in action-specific MEP modulation (Cohen’s d = 1.08) than simple attention, as tested in the counting task (Cohen’s d = 0.54). When the non-targeted grasp is observed, action-specific modulation is strongly suppressed (muscle p = .26) and no difference exists between the reward and the counting condition (task p = .37).

Follow-up analyses testing whether MEP_mod_ deviated significantly from zero revealed action-specific facilitation during the observation of rewarded target grasps in both FDI (p_FDR_ = .028, Cohen’s d = - 0.341) and ADM (p_FDR_ <.001, Cohen’s d = 0.717) muscles. During the observation of counted target grasps only the ADM muscle showed action-specific facilitation (p_FDR_ <.001, Cohen’s d = 0.628).

In summary, this pattern of MEP modulation indicates that action observation activates motor representations specific to different grip types in M1 and that this action specific neural tuning is significantly more pronounced when a high reward value is associated with the observed movement. This effect appears to be significantly larger than attention per se, as tested in the counting control condition.

## 3. Discussion

In this study, we showed that the value associated with observed actions significantly influences both behaviour and neural activity in human M1, and that these effects can be dissociated from attentional engagement. We show that highly valued observed actions lead to increased precision (decreased variability) of response times. Additionally, we show that there is stronger muscle-specificity of the physiological responses - suggesting “sharper” neural tuning - when highly valued actions are observed. Importantly, this putative strengthening of action-specific M1 response is not simply due to differences in the allocation of attention but rather to a more targeted effect of reward on motor resonance.

### 3.1. Reward influences behavioural responses to action observation

In our experiment, we recorded responses to observed actions in the context of high reward, low reward and a counting control condition and found that response times were least variable (i.e., lowest sigma values) in the high reward condition (**Figure 4**). Sigma characterizes the variability of the normal component of the RT distribution (Lacouture, 2008) so lowest sigma in the high reward condition indicates that the variability of the fast responses is lowest when value associated with the observed action is highest. Importantly response errors were low and mu (representing the mean RT) was similar across conditions reflecting that our task instructions explicitly favoured accuracy over speed and ensured that attentional engagement was high across conditions.

Our results are in line with previous work showing that RT variability was negatively correlated with motivation levels (Wang, Ding, & Kluger, 2014). Therefore, we can speculate that higher motivation levels in the high reward condition contributed to a narrowing of the RT distribution. Recent studies focusing on eye movements reported a similar improvement in motor precision with increasing reward value (Manohar et al., 2015; Manohar, Muhammed, Fallon, & Husain, 2019).

### 3.2. Action-specific representation of target versus non-target grasps in M1

We operationalized action-specific motor resonance as muscle-specific changes in corticomotor excitability as measured with single-pulse TMS simultaneously in the first dorsal interosseous (FDI) and the abductor digiti minimi (ADM) muscles when subjects observed whole-hand versus precision grip actions. This method offers the advantage of extracting muscle-specific information revealing the neural tuning of action representations evoked by movement observation (Alaerts, Senot, et al., 2010; Alaerts, Swinnen, & Wenderoth, 2010; Bunday et al., 2016; Cretu et al., 2019; de Beukelaar et al., 2016). Such a “two-action x two muscle” design is essential for dissociating the functional role of motor resonance from unspecific facilitation of the motor system (Cavallo et al., 2014) which is most likely caused by widespread general effects on the state of arousal during action observation experiments (Grafton, Fadiga, Arbib, & Rizzolatti, 1997; Murata et al., 1997; Valchev et al., 2015).

By quantifying the amount of action x muscle-specific facilitation in M1 via MEP_mod_ we showed a stronger neural tuning for target than non-target grasps when collapsed across reward and counting conditions. When the observed grasp was the non-target action (i.e., the action is not rewarded and not counted), action-specific modulation was strongly suppressed. Importantly, in our experiment, this suppression does not involve a general decrease in corticomotor excitability during non-target grasps. When collapsing across grasp types (i.e., PG and WHG), no differences were observed in the amplitudes of MEPs recorded during target and non-target trials in the two muscles, as shown in Supplementary Figure 1. A similar facilitation of task-relevant items was found in previous studies and was attributed to differences in selective attention (Banich et al., 2001; Egner & Hirsch, 2005; Gál et al., 2009; Polk, Drake, Jonides, Smith, & Smith, 2008).

### 3.3. Reward enhances the neural tuning of action representations in M1 during action observation over and beyond attentional effects

We show that monetary rewards associated with observed actions strengthen action-specific motor resonance (Cohen’s d = 1.09) significantly beyond the modulation already induced by task-relevant attention-driven salience (Cohen’s d = 0.54). This action-specific increase in motor resonance was overall stronger in the ADM muscle during the observation of both reward (Cohen’s d = 0.717) and counting (Cohen’s d = 0.628) target grasps. In the FDI muscle, action-specific modulation was only visible when the target action was rewarded (Cohen’s d = −0.341). Similar weaker patterns of action-specificity in the FDI muscle have been observed in previous research and attributed to the fact that FDI contributes to both WHG and PG, whereas ADM is involved specifically in the formation of WHG (Alaerts et al., 2012; Bunday et al., 2016; Davare, Lemon, & Olivier, 2008; Sartori, Betti, & Castiello, 2013). Note that for the target grasps, reward and counting conditions were closely matched, since participants had to count either the accumulated rewards or the occurrence of the target grasp while the overall protocol as well as the action stimuli were identical.

Although it is difficult to strictly dissociate attention and reward-related effects, we argue that the strengthening of action-specific signals in M1 during movement observation is mainly driven by the latter. In essence, our results suggest that reward modulates motor resonance during movement observation through the amplification of action specific representations in M1 (**Figure 5**).

Our findings regarding the modulation of motor resonance by action value (i.e., in the reward versus counting target conditions) are in good agreement with previous results from single neuron recordings in area F5 of primate premotor cortex (Caggiano et al., 2012). Caggiano and colleagues suggested that mirror neurons are modulated by the subjective value of observed actions, with the majority discharging more strongly during the observation of highly valued actions. Here, we investigated the modulation of action-specific motor resonance elicited in the human M1 during the observation of rewarding actions. Despite the obvious methodological differences between the studies, we complement the single cell results in monkeys by showing a similar strengthening of M1 mirroring during the observation of highly-valued actions in humans. The general assumption is that M1 motor resonance is driven by mirror neuron activity in up-stream areas, most notably ventral premotor cortex (de Beukelaar et al., 2016; Kilner, Neal, Weiskopf, Friston, & Frith, 2009). It is worth mentioning that in the experiment by Caggiano and colleagues, an actor grasped objects which differed with respect to their subjective value for the monkey (e.g. valuable food items versus objects with no value) and which were directly offered to the monkey on a subset of trials. By contrast, in our experiments, the value was directly associated with a specific grasp type at the start of each block, while the grasped item was identical. Nevertheless, similar to the design of Caggiano and colleagues we compared signals elicited during the observation of actions associated with reward or not, while levels of attention were similar across conditions. Therefore, our results extend these non-human primate findings by showing that action value influence motor resonance in humans over and beyond attentional effects.

Our findings are also in line with recent insights into the mechanisms of reward-based modulation which support an improvement of the signal-to-noise ratio of task-relevant characteristics (e.g., Engelmann and Pessoa, 2007; Etzel et al., 2008; Baldassi and Simoncini, 2011; Chelazzi et al., 2013; Grueschow et al., 2015; Zhang et al., 2018). Etzel and colleagues (2016) used pattern classification methods and showed that task rules were decoded more effectively on reward trials. In agreement to our findings, they proved that reward does not only lead to a generalized increase in brain activity but rather an enhancement of task-relevant representations.

Even though the enhancement of action specific tuning in M1 was larger for reward than for attentional engagement alone, it is unclear whether these net effects were driven by distinct neural mechanisms. Attending to task-relevant features can lead to a selective activity increase in neurons that encode those features in sensory areas (Maunsell & Cook, 2002). Reward can cause similar effects and it has been argued that it might impact on the strategic control of attention such that the prospect of earning a larger reward biases attention towards task-relevant stimuli in order to maximize overall gain (Chelazzi and colleagues, (2013). This is in agreement with recent findings suggesting that when value is associated with specific stimuli, these will subsequently have increased attentional priority (Anderson, 2013; Hickey, Chelazzi, & Theeuwes, 2010). Our results are principally in line with this proposal, however, it is important to note that reward specific effects were probed in the motor system. Several studies observed that M1 excitability is modulated by direct rewards or by reward expectation during motor tasks (Bundt, Abrahamse, Braem, Brass, & Notebaert, 2016; Gupta & Aron, 2011; Kapogiannis et al., 2008; Klein-Flugge & Bestmann, 2012; Mooshagian, Keisler, Zimmermann, Schweickert, & Wassermann, 2015; Suzuki, Hamaguchi, & Matsunaga, 2018; Suzuki et al., 2014; Thabit et al., 2011). However, showing that reward influences action specific representation in M1 even if no motor response is required is a new finding.

It is currently unclear at which stage reward information is integrated into the perceptual-to-motor transformation of action stimuli. Based on previous research, it is likely that PMv might be one important area mediating this effect. First, Caggiano et al., (2012) found a similar value-based modulation in PMv mirror neurons as observed here. Second, action specific PMv-M1 interactions have previously been demonstrated for both performing and observing precision versus whole hand grasping movements (Davare et al., 2008; Davare, Montague, Olivier, Rothwell, & Lemon, 2009; de Beukelaar et al., 2016; Fluet, Baumann, & Scherberger, 2010). Our results are consistent with the idea that reward optimizes grasp-specific PMv-M1 interactions, resulting in the observed sharpening of neural action representations in M1.

## 4. Acknowledgements

This work was supported by the SNSF Research Grant 320030_175616.

