## Supplementary Figures for "Reward modulates behaviour and neural responses in motor cortex during action observation"

**Supplementary Materials**

General corticomotor excitability changes

We investigated whether any general corticomotor excitability differences exist during the observation of target versus non-target grasps. We did this by calculating the average MEP amplitude during the grasp observation (i.e. collapsing across PG and WHG) separately for each muscle (i.e., FDI and ADM), condition (i.e., reward, count) and target (i.e., target, non-target). We performed a repeated-measures ANOVA with MEP amplitude as dependent variable and *muscle* (FDI, ADM), *task* (reward, count) and target (target, non-target) as within-subject factors.

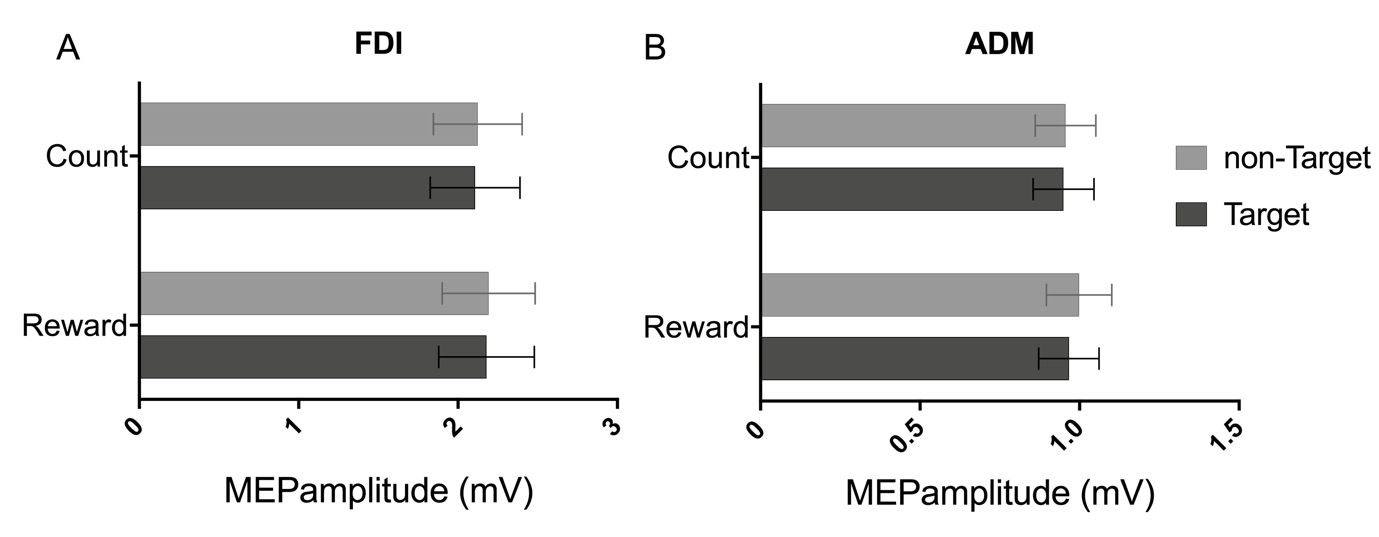
Results showed that general MEP amplitudes were higher in the FDI compared to ADM muscle (main effect of muscle p < 0.001) as well as in the reward condition compared to the counting task (main effect of task p=0.038). We did not find a main effect or interactions with the factor target (all p >=0.194). These results suggest that non-specific corticomotor changes during grasp observation (collapsed across grasp types) do not differ depending on the attentional demands of the task (supplementary figure 1).

**Supplementary Figure 1.** Motor- evoked potential (MEP) amplitude during the observation of grasping movements (average of whole hand grip and precision grip; n=34). Results are presented separately for each muscle [first dorsal interosseous (FDI) and abductor digiti minimi (ADM)], task (reward, counting) and target (target: dark gray; non-target: light gray).

Individual muscle-specific corticomotor changes

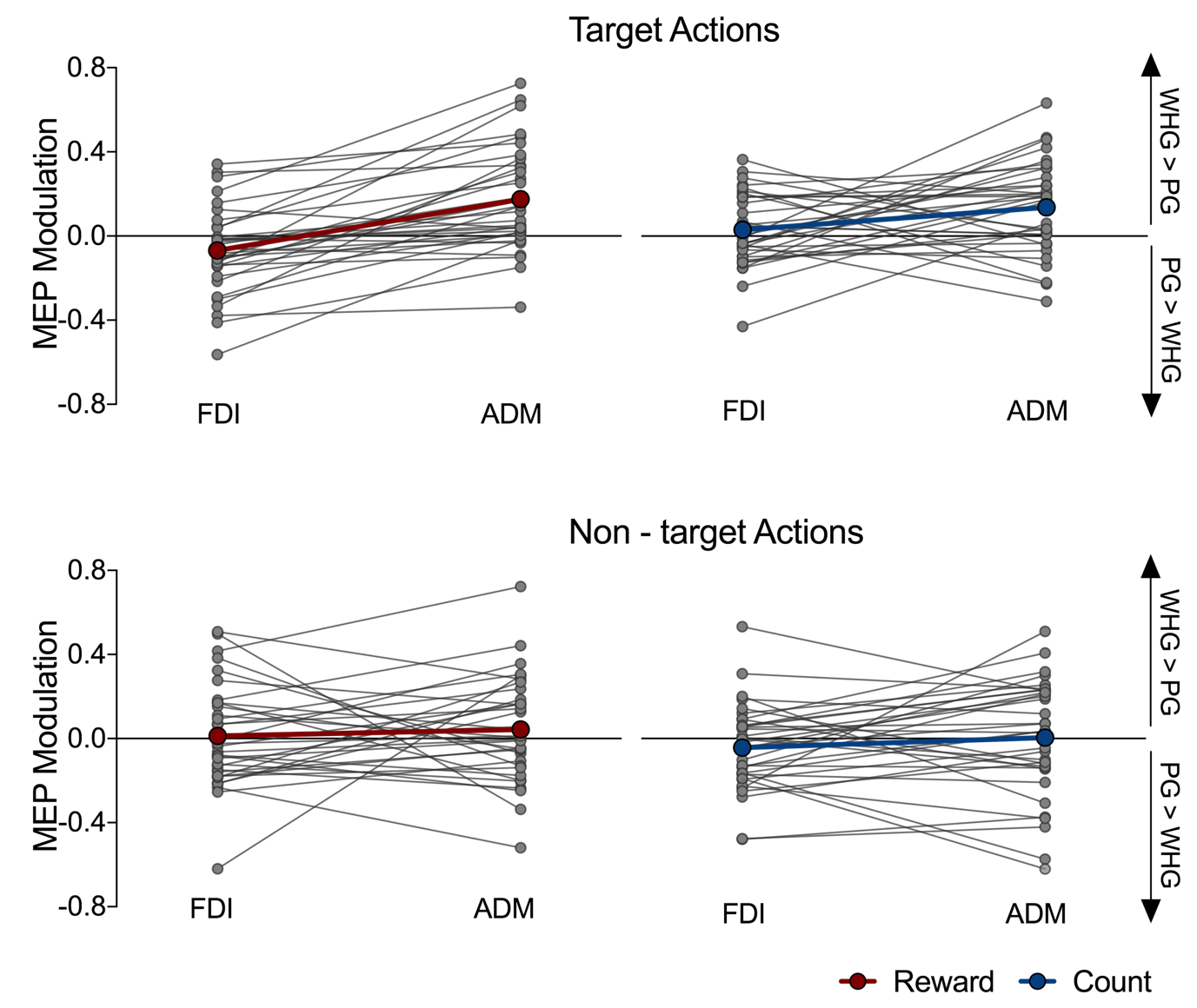

**Supplementary Figure 2.** Motor- evoked potential (MEP) modulation recorded in individual subjects during the observation of videos depicting a whole hand grip (WHG) or a precision grip (PG) either in a reward (red lines) or counting (blue lines) task. Results are presented separately for each target type and muscle [first dorsal interosseous (FDI) and abductor digiti minimi (ADM)]. Values >0 indicate higher facilitation during WHG vs. PG observation, whereas values <0 indicate increased facilitation during PG vs. WHG observation.

**Supplementary Table 1. Descriptive statistics** (mean ± standard deviation) for the MEP amplitude recorded in the TMS experiment. Results are presented separately for each condition and muscle [first dorsal interosseous (FDI) and abductor digiti minimi (ADM)].

|  | **FDI muscle** | | | |
| --- | --- | --- | --- | --- |
|  | **Target reward** | **Target count** | **Non-target reward** | **Non-target count** |
| **WHG** | 2.126 ± 1.736 | 2.144 ± 1.700 | 2.196 ± 1.726 | 2.049 ± 1.525 |
| **PG** | 2.231 ± 1.797 | 2.072 ± 1.616 | 2.187 ± 1.719 | 2.199 ± 1.760 |
|  | **ADM muscle** | | | |
| **WHG** | 1.065 ± 0.630 | 1.030 ± 0.664 | 1.006 ± 0.609 | 0.964 ± 0.578 |
| **PG** | 0.867 ± 0.501 | 0.869 ± 0.487 | 0.991 ± 0.613 | 0.948 ± 0.562 |
